# Improved Search of Large Transcriptomic Sequencing Databases Using Split Sequence Bloom Trees

**DOI:** 10.1101/086561

**Authors:** Brad Solomon, Carl Kingsford

## Abstract

Enormous databases of short-read RNA-seq sequencing experiments such as the NIH Sequencing Read Archive (SRA) are now available. These databases could answer many questions about the condition-specific expression or population variation, and this resource is only going to grow over time. However, these collections remain difficult to use due to the inability to search for a particular expressed sequence. While some progress has been made on this problem, it is still not feasible to search collections of hundreds of terabytes of short-read sequencing experiments. We introduce an indexing scheme called Split Sequence Bloom Tree (SSBT) to support sequence-based querying of terabyte-scale collections of thousands of short-read sequencing experiments. SSBT is an improvement over the SBT [1] data structure for the same task. We apply SSBT to the problem of finding conditions under which query transcripts are expressed. Our experiments are conducted on a set of 2,652 publicly available RNA-seq experiments contained in the NIH for the breast, blood, and brain tissues. We demonstrate that this SSBT index can be queried for a 1000 nt sequence in under 4 minutes using a single thread and can be stored in just 39 GB, a five-fold improvement in search and storage costs compared to SBT. We further report that SSBT can be further optimized by pre-loading the entire index to accomplish the same search in 30 seconds.

## 1 Introduction

An enormous amount of DNA and RNA short read sequence data has been published world wide. The NIH Sequence Read Archive (SRA) [2] alone contains almost 4 petabases of open-access sequence and continues to grow at an accelerating rate. This collection could be a great resource for understanding genetic variation, and condition- and disease-specific gene function in ways the original depositors of the data did not anticipate. For example, a natural use would be to search all public, human RNA-seq short-read files in the SRA (representing individual sequencing runs) for the presence of a particular transcript of interest to understand conditions of expression or develop a manageable subset for further analysis. However, searching the entirety of such a database for a query sequence has not been possible in reasonable computational time.

Some progress has been made toward enabling sequence search on large databases. The NIH SRA does provide a sequence search functionality [3]; however, it requires the selection of a small number of experiments to which to restrict the search. Existing full-text indexing data structures such as Burrows-Wheeler transform [4], FM-index [5], or others [6, 7, 8] cannot at present be efficiently built at this scale.Word-based indices [9, 10], such as those used by Internet search engines, are not appropriate for edit-distance-based biological sequence search. The sequence-specific solutions caBLAST and its variants [11, 12, 13] require an index of a known genomes, genes, or proteins and so cannot search for novel sequences in unassembled read sets. Further, all of these existing approaches do not handle the additional complication that a match to a query sequence *q* may span many short reads.

More recently two methods have been developed that store kmer approximations of experiments in directly searchable indices. The Sequence Bloom Tree [1] encodes all kmers in an experiment into a single bloom filter and builds a directly searchable binary tree of bloom filters over increasingly large subsets of the data. Queries are processed by looking up each kmer in a query for their presence or absence in the tree and recursing until all matching leaves have been found. It represents the current best method for searching a large database but cannot handle petabase-scale data. The Bloom Filter Trie [14] was designed as a direct compression method for a pan-genome but also can be queried for a set of kmers.

We address the search problem of finding all the experiments that contain enough reads matching a given query sequence *q* to indicate that *q* was present in the experiment. A query is an arbitrary sequence such as a transcript, and we find the collection of short-read experiments in which that sequence is present. These estimates themselves could be used to analyze conditions of gene expression or could make downstream analysis more efficient by filtering a large database for the relevant files. The Sequence Bloom Tree (SBT) was the first data structure to directly address this problem and could search a 5 terabase dataset in under 20 minutes using a 200 GB index. We modify the base structure of SBT with a new indexing data structure called Split Sequence Bloom Tree (SSBT). Like SBT, SSBT is independent of the eventual queries, so the approach is not limited to searching only for known genes, but can potentially identify arbitrary sequences. SSBTs can be efficiently built, extended, and stored in limited space and do not require retaining the original sequence files. Using SSBT, datasets can be searched using low memory for the existence of arbitrary query sequences. We compared SSBT against BFT and SBT and found that it outperforms both in terms of query time (5 times faster than SBT and 12 times faster than BFT) and storage cost, at the price of some additional time and temporary storage needed to construct the index.

## 2 Methods

### 2.1 Split Sequence Bloom Tree

The main idea behind SSBT is the creation of a hierarchy of compressed bloom filters [15, 16], which is used to efficiently store a set of experiments each consisting of many short reads (Figure 1). A bloom filter is a probabilistic storage structure which encodes an arbitrary set into a fixed length bit vector using hash functions. Here, each bloom filter encodes set of *k*-mers (length-*k* subsequences) present within a subset of the sequencing experiments and are stored using hash functions that convert these kmers to a specific index on the filter. We define each experiments’ individual kmer content as the set *b_i_*, with the collection *B* = {*b_i_* | 0 ≤ *i* < *n*} denoting a set of *n* experiments represented by their kmer content. Throughout, we abuse notation slightly to identify bloom filters with the sets they represent.

**Figure 1:**
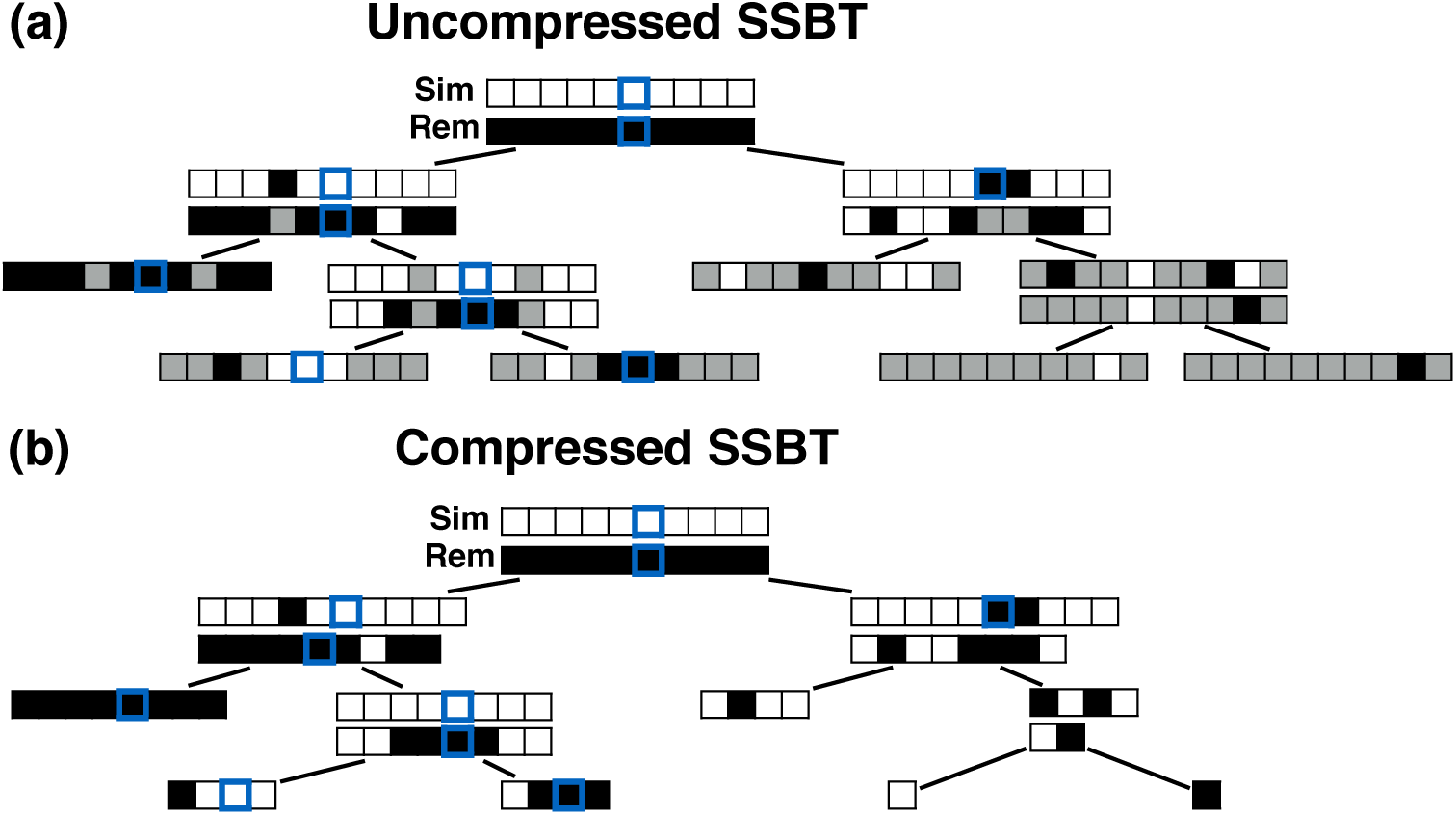
An example uncompressed and compressed SSBT where black corresponds to a bit value of ‘1’ and white corresponds to a bit value of ‘0’. (a) Grey bits correspond to non-informative bits whose value is known given a parent filter. We see that grey bits are cumulative and exist at all index positions below a on ‘1’ in the sim filter or a ‘0’ in the dif filter. When looking up index value 6, each filter is queried until either a sim ‘1’ is found or a dif ‘0’ is found. (b) All non-informative bits have been removed from the uncompressed tree. The lookup for index value 6 is adjusted based on the removed non-informative bits.

An SSBT is a binary tree that stores each *b_i_* in *B* across a unique path from root to leaf with each leaf mapping to a single experiment. The root node *r* of an SSBT contains the total content of each *b_i_* and stores this information using two identically sized bloom filters with a single conserved hash, the pair of which we define as a single ‘split bloom filter’. We define the first filter as the *similarity filter* 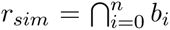 as the bloom filter which stores the universally expressed kmers in set *B*. The second filter is defined as the *remainder filter* 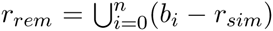 as the bloom filter which stores the remaining kmers — those in the union of all *b_i_* excluding those found in the similarity vector. By this definition, the bitwise union of *r_sim_* and *r_rem_* is equivalent to a single fixed length bloom filter which stores all *b_i_* in *B*. Furthermore, the bitwise intersection of *r_sim_* and *r_rem_* is the nullset. Different kmers in *B* may hash to the same position in *r_sim_*. As *r_rem_* is defined by *r_sim_*, additional ‘similarity’ may be found by random chance when hashing kmers. The inverse (identical kmers mapping to different positions) is not possible in a bloom filter.

Given the root *r*, only the kmers which are stored in *r_rem_* are then passed to the children. In this way *r*’s immediate children do not store the full set *b_i_* but rather the modified 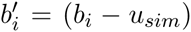. By repeating this down the path from root to leaf, the set of kmers being stored depletes gradually until a leaf is found.This leaf stores just the remaining kmers in 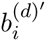, where *d* is the depth of the leaf. As the intersection of a single set is the set itself, the leaf’s sim filter is the set 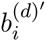 and the leaf’s rem filter is the 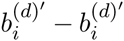, or the nullset. We can recover the original set by following unique path from root to leaf and find 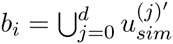, where *d* is the length of the path to leaf repository *i*.

### 2.2 SSBT Construction and Insertion

A SSBT is built by repeated insertion of sequencing experiments followed by the removal of all non-informative bit indices from each filter. Given a (possibly empty) SSBT *T*, a new sequencing experiment *s* can be inserted into *T* by first computing the fixed-length bloom filter *b_s_* of the kmers present in *s* and then walking from the root along a path to the leaves and inserting *s* at the bottom of *T* in the following way. When at node *u*, if *u* has children, *b_s_* has to be split between *u_sim_* and *u_dif_*. This is done through the bit updates defined by Table 1 for each bit index *i* in 0 ≤ *i* ≤ |*b_s_*|. These updates ensure that *u* correctly stores what is still universally similar in *u_sim_* and what now exists below *u* in the tree with *u_dif_* and that *b_s_* has been updated to store similar elements at *u*.

**Table 1:** Bit update table when inserting *b*(*s*) into the subtree rooted at *u*. *u*’s two immediate children are both updated with the single column *c_sim_*. A value of ‘-’ implies that no change needs to be made to that bit. (i) *b*(*s*) contains a value already stored in *u_sim_*, the value is removed from *b*(*s*). (ii) *b*(*s*) does not contain a value stored in *u_sim_*. Although the bit is no longer similar at *u*, *b*(*s*) has not yet been inserted into a child and all current children of *u* are universally similar at this location. (v) *b*(*s*) contains a value not found in the tree. *u_rem_* is updated to reflect that the value now exists. (iii, iv, vi) No changes need to be made.

|  | Before |  |  | After |  |  |  |
| --- | --- | --- | --- | --- | --- | --- | --- |
| | $u_{sim}$ | $u_{rem}$ | $b(s)$ | $u_{sim}$ | $u_{rem}$ | $b(s)$ | $c_{sim}$ |
| (i) | 1 | 0 | 1 | - | - | 0 | - |
| (ii) | 1 | 0 | 0 | 0 | 1 | - | 1 |
| (iii) | 0 | 1 | 1 | - | - | - | - |
| (iv) | 0 | 1 | 0 | - | - | - | - |
| (v) | 0 | 0 | 1 | - | 1 | - | - |
| (vi) | 0 | 0 | 0 | - | - | - | - |

The potentially modified *b_s_* is then compared against each child *c_i_* of *u* in order to find the most similar child. While we tested many similarity metrics for comparing a single filter *b*(*s*) with a split filter (*c_i,sim_*, *c_i,dif_*), the metric used in this manuscript first computes the total number of 1-bits shared between the *c_i,sim_* and *b_s_*. If no child has any 1-bits in common with *b_s_*, the metric then selects the child with the smallest Hamming distance between *b_s_* and *c_i,rem_*(*s*). Once the most similar child has been found, this child then becomes the new current node, and the process is repeated with a new subtree. If *u* has no children, then *u* represents a sequencing experiment *s*′ and only contains one vector *u_sim_*. In this case, a new node *v* is created as a child of *u*’s parent with *u* and *b_s_* as *v*’s children. As both *u* = *u_sim_* and *b_s_* are leaves, *v_sim_* is the bit-intersection of *u* and *b_s_* while *v_rem_* is the bit-union of *u* − *v_sim_* and *b_s_* − *v_sim_*. Finally we remove the elements in this new parent node from both children by replacing *u* with *u* − *v_sim_* and *b_s_* with *b_s_* − *v_sim_*. This yields an uncompressed SSBT containing all previous nodes and two new nodes *v* and *b_s_*.

### 2.3 Bit non-informativity in SSBT

Given the definition of SSBT’s split filters described above, for an arbitrary node *u* the only values allowed at a particular index *i* are (*u_sim_*[*i*], *u_rem_*[*i*]) = (1, 0), (0, 1), (0, 0). However, every index is represented using a bit from either filter, even when one filter’s value clearly defines the other. We address this inefficiency by removing these “non-informative bits” from the tree. Here we define a non-informative bit as a bit index *i* present in node *u* whose value can be determined by examining the bloom filters present in the set of parent filters above *u*. We describe the cases of non-informativity in SSBT below and provide an example in Figure 1:

1. For a bit index *i*, if *u_rem_*[*i*] = 0, then for every descendant *c* of *u*, *c_sim_*[*i*] = *c_rem_*[*i*] = 0 and *i* is non-informative below *u*. This is a direct result of the definition of *u_rem_* as the union of all children below it.
2. For a bit index *i*, if *u_sim_*[*i*] = 1 then *u_rem_*[*i*] = 0 and *u_rem_*[*i*] is non-informative. This is a formalization of the fact that the rem filter only contains the elements which are not stored in the sim filter.
3. For a bit index *i*, if *u_sim_*[*i*] = 1 then for every descendent *c* of *u*, *c_sim_*[*i*] = *c_rem_*[*i*] = 0 and *i* is non-informative below *u*. This is the logical extension of applying (1) to (2).

Using these cases, we can remove all non-informative bits from *u_rem_* given *u_sim_* and we can remove all non-informative bits from *u*’s immediate children using both *u_sim_* and *u_rem_*. As bits are only ever removed,for a node *u* and its child *c*, |*c_rem_*| ≤ |*c_sim_*| ≤ |*u_rem_*| ≤ |*u_sim_*|. We further note that the size reduction at each step are proportional to the similarity in experiments stored at *u*, with both uniform expression and non-expression of specific kmers lead to size reductions for all subsequent filters.

### 2.4 SSBT Compression

Given an uncompressed SSBT *T* with bloom filters of fixed length *m* and conserved hash function, we build a compressed SSBT *T*′ by removing the set of all non-informative bits (defined in Section 2.3) from *T* and compressing the variable length filters using RRR-compression [17]. We handle the ‘removal’ of a bit in node *u* by constructing an intermediate node *u*′ and copying only the informative bits from *u* to *u*′. The total number of informative bits in *u_sim_* and *u_rem_*, as well as their exact indices, can be determined using two filters: *u*’s immediate parent’s rem filter, *p*(*u*)*_rem_*, and *u_sim_*. Here we define *rank_x_*(*u*)[*j*] to be the number of bits set to *x* in the bit vector *u* from index 0 ≤ *i* < *j*. This can be computed in constant time using the C++ package sdsl [19].

Given *p*(*u*)*_rem_*, the only informative bits in *u_sim_* are those *i* for which *p*(*u*)*_rem_*[*i*] = 1 and 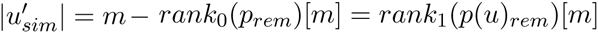. Likewise given *u_sim_*, the only informative bits in *u_rem_* are those *i* in which both *u_sim_*[*i*] = 0 and *p*(*u*)*_rem_*[*i*] = 1 and 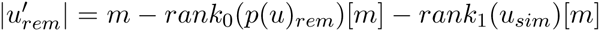. At each *i*, the values in *p*(*u*)*_rem_*[*i*] and *u_sim_*[*i*] determine if *i* is informative. If *i* is informative, it is copied over to the next open position in 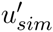 and/or 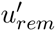. Subsequently, *u*′ is compressed via the RRR [17] compression scheme, which allows querying a bit without decompression and incurs only a *O*(log *m*) factor increase in access time (where *m* is the size of the bloom filter with non-informative bits removed). This process operates for every node in the tree, starting with the root node *T* which has a full length 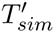 as it has no parent. See Figure 1 for an example of the compression step.

### 2.5 SSBT Querying

Given a query sequence *q* and a compressed Split Sequence Bloom Tree *T*, the sequencing experiments that contain *q* can be found by breaking *q* into its constituent set of kmers *K_q_* and then flowing these kmers over *T* starting from the root. In an SSBT, these kmers are organized into a set of unique kmers and immediately converted into a vector of filter indices *V_q_* using the conserved hash functions defined by *T*’s root’s sim filter. At the root node *u*, we query first *u_sim_* for each index in *V_q_*. Matches in *u_sim_* are recorded as ‘universal hits’ since they are found in intersection of all experiments rooted beneath *u*. Indices that are universal hits do not have to be queried further and are removed from the set — the sum of these hits records their presence at all children to *u*. The remaining indices that were not found in *u_sim_* are then queried in *u_rem_*. As |*u_sim_*| ≥ |*u_rem_*|, this query is accomplished by adjusting all indices *V_i_* ∈ *V_q_* to account for the non-informative bits removed. As we have already converted kmers to hash values, we transition from *u_sim_* to *u_rem_* by subtracting the number of 1-bits which occurred before *V_i_* in *u_sim_*. This is simply the *rank*_1_(*u_sim_*, *V_i_*), a property of a bit vector which can be computed in constant time using RRR-compressed vectors.

Each modified index can then be queried in *u_rem_*. Indices that map to 0-bits do not have to be queried further as they do not exist in any child to *u*. Indices that map to 1-bits are potential hits which belong to at least one child below *u* but not all. By adding the number of potential hits in *u_rem_* with the number of universal hits found in *u_sim_*, the number of matching kmers is determined for each query. If, for a user-specified cutoff *θ* ϵ [0, 1], this count is less than *θ* |*V_q_*|, then the query cannot exist in this subtree and the subtree is not searched further (it is pruned). If there exist *θ* |*V_q_*| or more universal matches, every child beneath *u* is a query hit and the tree also does not have to be searched further. Only in the case where the count is greater than *θ* |*V_q_*| but not enough universal matches have been found do we have to proceed to *u*’s children. To transition each index from node *u* to child node *c*, each index has to be further adjusted by the number of 0-bits in *u_rem_*. Once again, this can be calculated in constant time using *rank*_0_(*u_rem_*, *V_i_*). By repeating this process down through the tree, SSBT efficiently prunes branches that cannot contain the query, prunes queries which are known to exist in all children, and maintains a consistent hash function across a variable length set of compressed filters. An example of this query process can be found on Figure 1.

Using this logic, the set of index positions that define a query changes after each node is queried. To prevent having to store a unique query set for every node in the tree, we stored *V_q_* only once outside of the SSBT structure and developed a reversible means of activating or deactivating an index, as well as reversing changes made to the index value when descending the tree. Given a vector of indices *V_q_*, we define a single integer — the *tail-index* — to be the position along the vector that contains the last ‘active’ query index. This tail-index is initialized to the final value in *V_q_* and queries which are ‘removed’ simply swap positions with the tail-index and the tail-index is decremented by one. In such a way, we store the full set of indices but only query those indices up to and including the tail-index. By storing the tail-index present in each node (a cost of a single integer per node), we can restore all queries which were active at that node. Because the tail-index defines both the pivot between ‘active’ and ‘removed’ and the swap position for deactivating an index, the order in the vector records the order of deactivation. Because of this, any index which was ‘active’ for an arbitrary node *u* with tail-index *k* will always be among the first *k* indices in *V_q_*. Thus using the tail-index we can exactly store the unique set of indices that need to be searched at any node using only a single vector and an integer at each node.

In addition to activating or deactivating indices, the index positions themselves change between *u_sim_* and *u_rem_* and between *u* and *u*’s children. Given an index position *i_rem_* which maps to *u_rem_*, we can reconstruct the index position *i_sim_* which maps to *u_sim_* by looking up the *i_rem_*-th informative bit in *u_sim_*. As *u_sim_* = 1 defines non-informativity, we simply find the *i_rem_*-th 0 in *u_sim_*. For an RRR-compressed vector, this can be computed in *O*(log *m*) time for a length *m* vector using the *select*_0_(*u_sim_*)[*i_rem_*] operation. Likewise given an index position *j_sim_* mapping to an arbitrary child node of *u*, *c_sim_*, we can recover the index position *i_rem_* mapping to *u_rem_* by finding the *j_sim_*-th informative bit in *u_rem_*. As *u_dif_* = 0 defines non-informativity, we simply find the *j_sim_*-th 0 in *u_dif_*. For an RRR-compressed vector, this can be computed in *O*(log *m*) time for a length *m* vector using the *select*_1_(*u_dif_*)[*j_sim_*] operation. Thus, using just the rank and select operations implicit to an RRR-compressed vector, we can recover any index position at any node given a position along the SSBT and the SSBT split bloom filters themselves.

### 2.6 Accuracy

SSBT builds off of the base bloom filters used in SBT. For the same queries, it yields an identical set of results in a more efficient way. As it has been shown that kmer similarity is highly correlated to the quality of the alignments between sequences [21, 22, 23, 24], and SBT has previously determined the accuracy of this metric [1], we did not investigate the accuracy of SSBT, which is the same as SBT, further in this manuscript. In fact, one can think of SSBT as a lossless compression scheme on SBTs that operates before RRR-compression.

## 3 Results

### 3.1 Data and Hardware

All data used in this publication was constructed from the set or a subset of 2652 human, RNA-seq short-read sequencing runs from the NIH SRA [2]. At the time of download, these files consisted of the entire set of publicly available, human RNA-seq runs from blood, brain, and breast tissues stored at the SRA (as determined by keywords in their metadata and excluding files sequenced using the SOLID technology). This dataset has a total size of 5 TB of raw SRA data or 2.7 TB of gzipped fasta sequences. The 50 files selected for the comparison with BFT were chosen at random from this set and have a total size of 49 GB of gzipped fasta sequences. Kmers counts were computed using the Jellyfish 2.0 library for SBT and SSBT and KMC 2.0 for BFT. All times in these experiments were obtained on a shared computer with 16 Intel Xeon 2.60GHz cores using a single thread. BFT was run using default options with a compression constant of *k* = 5. SBT and SSBT use a kmer size of 20 while BFT was built using a kmer size of 18 as it only allows kmer lengths that are multiples of 9.

### 3.2 Evaluation on Build Time and Storage Cost

We compared the build time, maximum memory cost, and storage cost between SBT version 0.3.5, SSBT version 0.1, and BFT version 0.6. Both SBT and SSBT could be run on the full 2652 experiment dataset and both were run with a maximum of 100 tree nodes in memory. The results from this analysis are summarized in Table 2 and show that SSBT is roughly four times slower to build but yields a significantly smaller searchable index when compressed. As the indices only need to be build once (and can be incrementally build from the uncompressed state), the SSBT is a superior choice when there is sufficient hardware support for its larger uncompressed size. We further note that even this size (1853 GB) is significantly smaller than the raw data, that this size includes the bloom filter representation of every experiment, and that the raw data is not needed during the search once the bloom filter is constructed.

**Table 2:** Build statistics for SBT and SSBT constructed from a 2652 experiment set. The sizes are the total disk space required to store a bloom tree before or after compression. In SSBT’s case, this compression includes the removal of non-informative bits.

| Data Index | SBT | Split SBT |
| --- | --- | --- |
| Build Time | 18 Hr | 78 Hr |
| Compression Time | 17 Hr | 19 Hr |
| Uncompressed Size | 1295 GB | 1853 GB |
| Compressed Size | **200 GB** | **39.7 GB** |

We were unable to directly compare BFT with these numbers due to several complications encountered while running BFT. Specifically BFT version 0.6 has an unresolved error and frequently seg faults when inserting a large number of experiments. Because of this bug, we could not reliably produce a BFT with more than 50 total files. In order to compare BFT with the other existing methods, we built a separate index for BFT, SBT, and SSBT which contains only 50 experiments, randomly selected from the 2652-experiment set. For this small scale test, SBT and SSBT’s RAM loads were selected to be roughly equivalent to BFT. This was accomplished by setting SBT’s maximum in-memory filter load to 100 experiments and SSBT was limited to 30 experiments.

This is much smaller than the intended use case of SBT or SSBT but reflects the only fair comparison between tools. Even with this handicap, we see that BFT takes 16 times longer to build than the combined time to build and compress an SBT despite using more RAM. Similarly SSBT builds and compresses five times faster than BFT’s build and yields a directly searchable index with less than 1/9th the total storage cost. These results are summarized in Table 3 and Table 4. We note that BFT does have a smaller uncompressed size than either metric but that both SBT and SSBT were never meant to be queried using an uncompressed index and that the previously reported comparison in BFT’s publication did not properly use the necessary compression step.

**Table 3:** Build and compression peak RAM loads and on-disk storage costs for SBT, SSBT, and BFT constructed from a 50 experiment set. BFT does not have a built-in compression tool and cannot be queried when compressed. For these reasons, the uncompressed BFT is compared against the compressed SBT/SSBT.

| Data Index | BFT | SBT | SSBT |
| --- | --- | --- | --- |
| Build Peak RAM (GB) | 23 | 21.5 | 15.6 |
| Compress Peak RAM (GB) | - | 24.2 | 16.2 |
| Uncompressed Size (GB) | **9.2** | 24 | 35 |
| Compressed Size (GB) | - | **3.9** | **0.94** |

**Table 4:** Build and compression times for SBT, SSBT, and BFT constructed from a 50 experiment set. As SBT and SSBT were designed to be queried from a compressed state, we compare the time to build and compress against BFT’s time to build.

| Data Index | BFT | SBT | SSBT |
| --- | --- | --- | --- |
| Build Time (Min) | 195 | 6 | 19 |
| Compression Time (Min) | - | 6.5 | 17 |
| Total Time (Min) | **195** | **12.5** | **36** |

### 3.3 Evaluation on the Query Time

We evaluated the efficiency of queries in SBT, SSBT, and BFT on three sets of 100 queries. To build each query set, we estimated the expression profiles of all 50 experiments used in the small-scale indices using Sailfish [25]. We then randomly sampled transcripts that were expressed at TPM values at or above 100, 500, or 1,000 in at least one of those files to build three query sets, of 100 queries each. Each query was run individually for each tool and the file system cache was emptied at the end of each run to ensure that the average time is an accurate representation of query behavior. The results are summarized in Table 5 and show that SSBT is anywhere from 3–13x faster than either method at this scale. Although this is a significant improvement, we suspect that this 50-experimenttest underestimates SSBTs relative performance, due to SSBTs efficient storage of similar elements and better optimized querying.

**Table 5:** Comparison in query timing (and average peak memory) between SBT, SSBT, and BFT indices for 50 experiments.

| **Index** | **TPM $\geq 100$** | **TPM $\geq 500$** | **TPM $\geq 1000$** |
| --- | --- | --- | --- |
| BFT | 75 Sec (11.8 GB) | 75 Sec (11.8 GB) | 75 Sec (11.8 GB) |
| SBT | 19 Sec (2.9 GB) | 21 Sec (3.1 GB) | 22 Sec (3.2 GB) |
| SSBT | 5.8 Sec (0.64 GB) | 6.2 Sec (0.65 GB) | 6.3 Sec (0.66 GB) |

A better comparison was possible using the full 2652-experiment indices with SBT and SSBT. The query sets used in this analysis were randomly selected to exist in at least one of 100 randomly sampled experiments out of the full dataset with three TPM-specific sets constructed as before. The results are summarized in Table 6 and Table 7 and show that SSBT is over five times faster than SBT regardless of the TPM value or cutoff threshold used in either index.

**Table 6:**
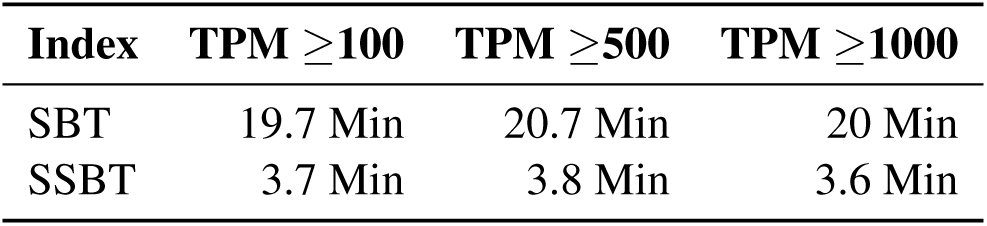
Comparison in query timing between SBT and SSBT for 2652 experiments.

Given that SSBT’s speed improvement closely mirrors its size improvement (a five-fold speedup for a five-fold size reduction), we hypothesized that SSBT could be made significantly faster by reducing or eliminating the I/O costs associated with loading and unloading bloom filters. This is only possible for an SSBT, whose directly searchable index using less than 1% of the size of the original data. This resulted in an additional 7x speedup over regular SSBT and a roughly 39x increase over SBT. We report this result as ‘RAM SSBT’ in Table 7.

**Table 7:** Comparison of query times using different thresholds *θ* for SBT and SSBT using the set of data at TPM 100.

| **Query Time:** | **$\theta=0.7$** | **$\theta=0.8$** | **$\theta=0.9$** |
| --- | --- | --- | --- |
| SBT | 20 Min | 19 Min | 17 Min |
| SSBT | 3.7 Min | 3.5 Min | 3.2 Min |
| RAM SSBT | 31 Sec | 29 Sec | 26 Sec |

SSBT’s speed improvement can generally be explained by a reduction in I/O costs but SSBT has another key benefit in the ability to prune queries which are found in every child (“universal query pruning”). This is not relevant for the average query but is a significant improvement in recovery of queries which are expressed in a large fraction of the database. We demonstrate this property by recording the number of SSBT nodes loaded in our TPM 100 set. When universal query pruning is ignored (Figure 2), queries that are expressed in a majority of the dataset are inefficient to look up. However, when query pruning is introduced (Figure 3), significantly fewer queries look at more than 2652 nodes.

**Figure 2:**
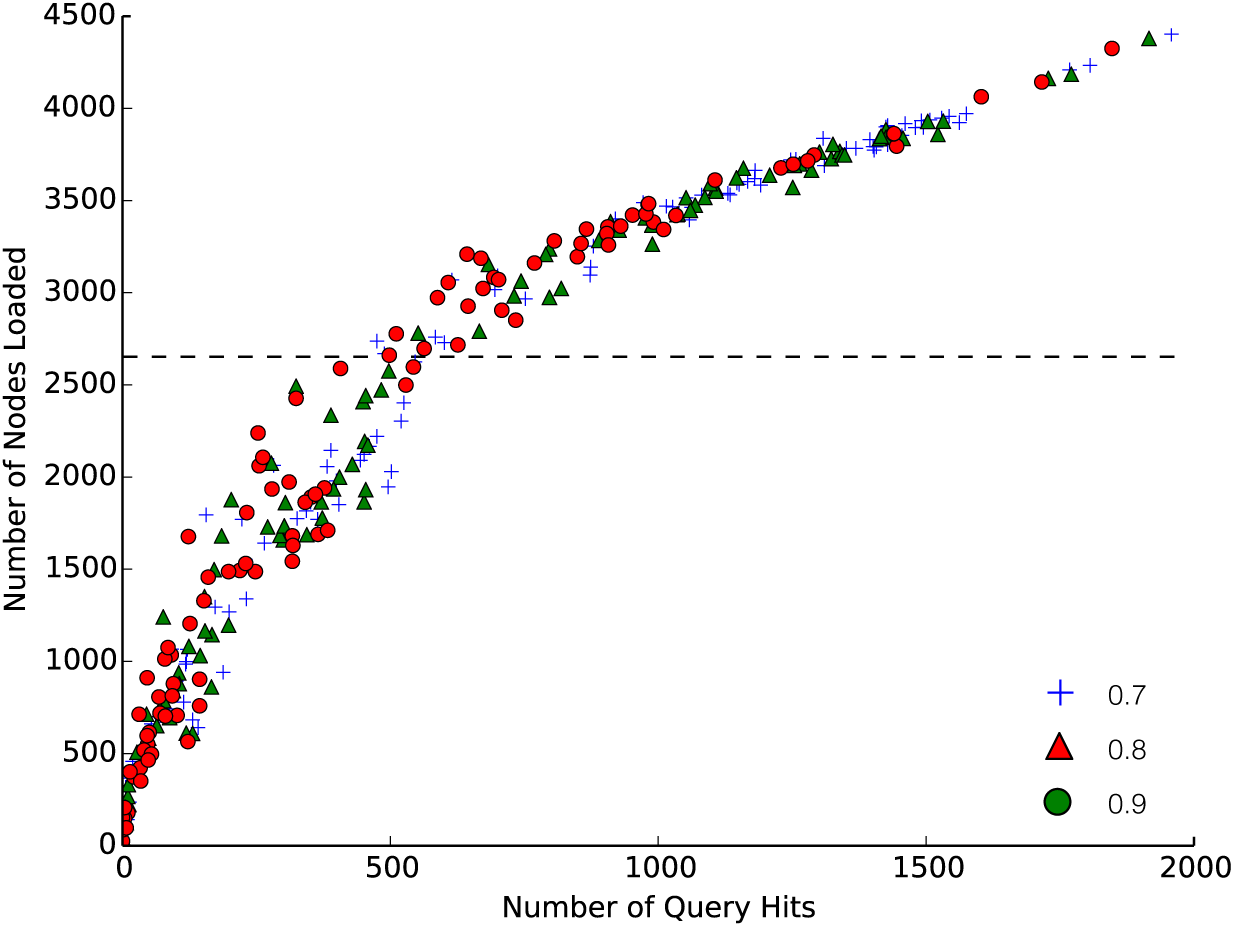
Number of SSBT nodes that would be loaded if SSBT did not prune queries against the total number of query matches found among 2652 experiments. Blue, green, and red correspond to a kmer matching threshold of 0.7, 0.8, and 0.9 respectively. A naïve approach would search all 2652 leaves, represented by the black dashed line.

**Figure 3:**
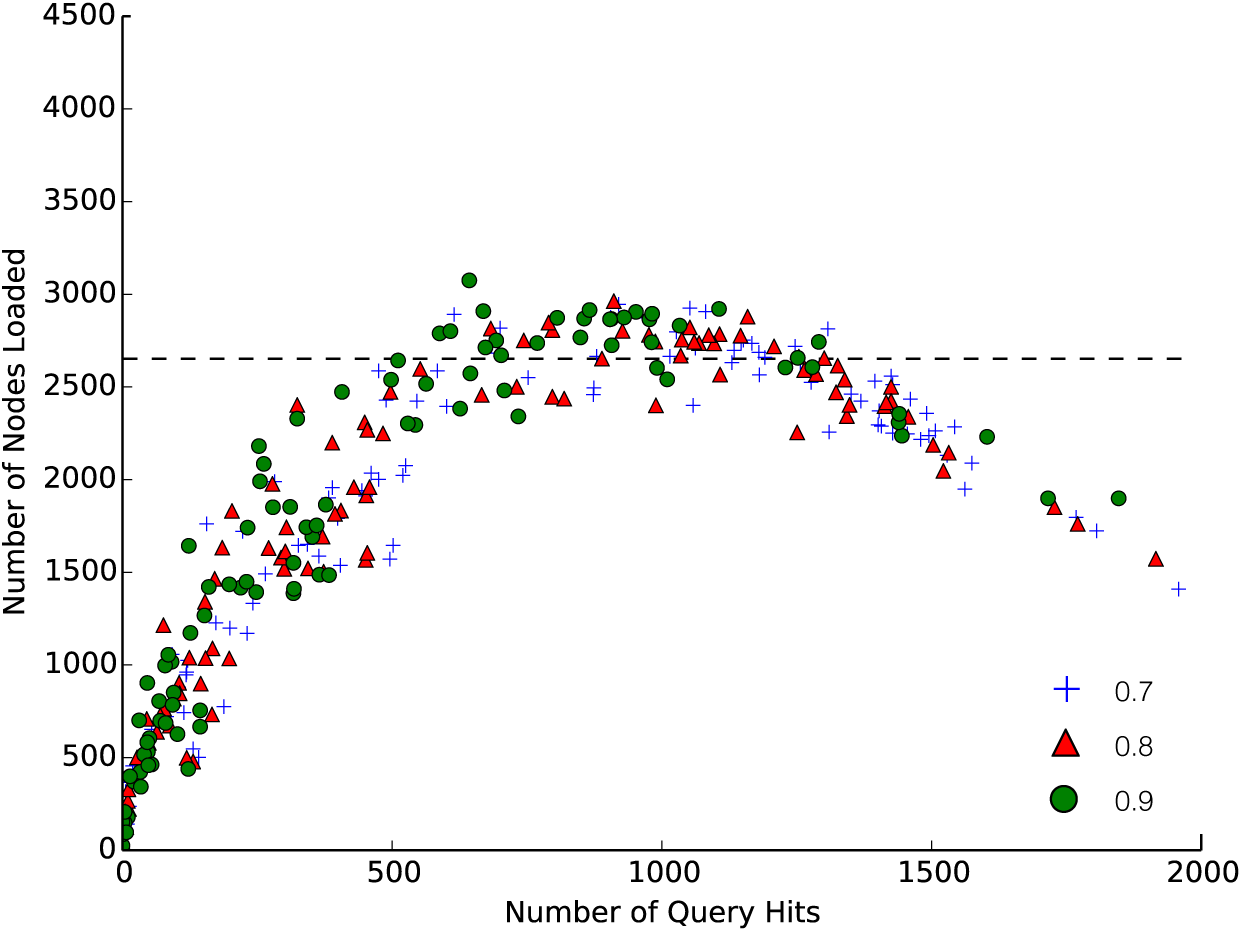
Number of SSBT nodes loaded against the total number of query matches found among 2652 experiments. Blue, green, and red correspond to a kmer matching threshold of 0.7, 0.8, and 0.9 respectively. A naïve approach would search all 2652 leaves, represented by the black dashed line.

## 4 Conclusion

The Split Sequence Bloom Tree is a novel approach to searching for short read experiments in a large database. It uses a more efficient encoding scheme to generate a compressed but directly searchable index which is at least five times smaller than any existing method. This improvement is significant for all queries but produces the largest gap over existing techniques when querying transcripts which are found in many experiments. SSBT’s improved storage allows 5 TB of sequencing information to be indexed in 40 GB, yielding a 5x increase in speed. Its on-disk memory usage scales more efficiently than any previous tool and is the only tool whose size permits the entire index to be loaded completely into RAM. As the query times of SSBT are bottlenecked by I/O, pre-loading an SSBT yields a 39x increase in speed over the closest competitor. Although these improvements come at a significant cost in build time and some additional uncompressed storage usage, these operations are typically much more rare than queries. All of the results in this paper were run using a single thread on a single computer. Future work optimizing SSBT for multiple-threaded builds and querying should produce an even more significant improvement in build and query times.

SSBT is open source and available at http://www.cs.cmu.edu/∼ckingsf/software/bloomtree/.

## Acknowledgements

We would like to thank Hao Wang, Natalie Sauerwald, Cong Ma, Tim Wall, Mingfu Sho, and especially Guillaume Marçais, Dan DeBlasio, and Heewook Lee for valuable discussions and comments on the manuscript. This research is funded in part by the Gordon and Betty Moore Foundation’s Data-Driven Discovery Initiative through Grant GBMF4554 to Carl Kingsford. It is partially funded by the US National Science Foundation (CCF-1256087, CCF-1319998) and the US National Institutes of Health (R21HG006913, R01HG007104). C.K. received support as an Alfred P. Sloan Research Fellow. B.S. is a predoctoral trainee supported by US National Institutes of Health training grant T32 EB009403 as part of the Howard Hughes Medical Institute (HHMI)-National Institute of Biomedical Imaging and Bioengineering (NIBIB) Interfaces Initiative.

